# Sugar distributions on gangliosides guide the formation and stability of amyloid-β oligomers

**DOI:** 10.1101/2023.05.09.540003

**Authors:** Jhinuk Saha, Brea J. Ford, Sydney Boyd, Vijayaraghavan Rangachari

## Abstract

Aggregation of Aβ peptides has been known as a key contributor to the etiology of Alzheimer’s disease. Being intrinsically disordered, the monomeric Aβ is susceptible to conformational excursions, especially in the presence of key interacting partners such as membrane lipids, to adopt specific aggregation pathways. Furthermore, key components such as gangliosides in membranes and lipid rafts are known to play important roles in the adoption of pathways and the generation of discrete neurotoxic oligomers. Yet, what roles the carbohydrates on gangliosides play in this process remains unknown. Here, using GM1, GM3, and GD3 ganglioside micelles as models, we show that the sugar distributions and cationic amino acids within Aβ N-terminal region modulate oligomerization of Aβ temporally, and dictate the stability and maturation of oligomers.

## 1. Introduction

Alzheimer’s disease (AD) is the most prevalent neurodegenerative pathology affecting elderly patients over 60 years of age across the world. The extracellular plaque deposits and the intracellular neurofibrillary tangles (NFTs) mainly comprised of amyloid-β (Aβ) peptides and hyperphosphorylated tau, respectively form the hallmark lesions in AD brains [1–5]. One of the major components of these Aβ plaques is Aβ42, a 42-amino acid long isoform β, aggregates of which are highly neurotoxic and contribute to synaptic dysfunction and memory impairment [6–9]. Aβ42 (hereon referred to as Aβ) undergoes self-assembly from the monomeric form to high molecular weight aggregates called protofibrils and fibrils and along the pathway, the low molecular weight soluble oligomers are known to be the primary toxic species involved in synaptic dysfunction and memory loss[8,10–24]. Aβ oligomers can be highly heterogeneous both in size and structure depending on the pathway of aggregation as well as the interacting partners [25–32].

Key cellular components that affect Aβ aggregation pathways and generation of oligomers are the membrane lipids[33–39]. Being a part of transmembrane amyloid precursor protein (APP), Aβ has a strong affinity for cellular membrane lipids[40], which has been well-established to influence Aβ aggregation [41–46]. For example, results from model liposomal systems with small unilamellar vesicles (SUVs) and large unilamellar vesicles (LUVs) have established that morphologically distinct Aβ42 fibrils can be generated under the influence of liposomal surfaces [38,47–57]. Furthermore, lipid components within the membranes and lipid rafts such as cholesterol and gangliosides have been known to modulate Aβ aggregation[39,58–66]. Particularly, membranes containing GM1 gangliosides induce the formation of conformationally distinct and cytotoxic Aβ fibrils[67,68]. Not only GM1-containing micelles and liposomes but free, non-micellar GM1 gangliosides too modulate Aβ aggregation pathways[69].

Furthermore, in addition to GM1, other gangliosides present within the membranes of different cell types[70–73] selectively interact with Aβ and its mutants[71–73], and influence aggregation kinetics [71,72,74–76]. We have also previously shown that GM1 ganglioside micelles induce the formation of oligomers with distinct biophysical and biochemical properties[26]. However, it remains unclear as to how sugar distributions on gangliosides confer specificity of interactions with Aβ and what specific residues contribute to these interactions. To address these questions, here we chose three of the key gangliosides with different sugar distributions present in membrane microdomains of neuronal cells, smooth muscles, and neural stem cells, i.e., GM1, GM3, and GD3 micelles to investigate their effect on oligomerization of Aβ42. Using three specific mutants of Aβ42; R5A, K16A, and H^13^H^14^AA double mutant and along with wildtype, here we uncover that sugar distributions are key in guiding Aβ oligomerization and fibrillation along what seems to be different pathways.

## 2. Materials and methods

Columns for size exclusion chromatography (SEC) (Superdex-75 HR 10/30) were purchased from Cytiva (Marlborough, MA). Zorbax C8 HPLC column was bought from Agilent technologies (Santa Clara, CA). GM1, GM3 and GD3 gangliosides were purchased Avanti Polar Lipids, Inc. (Alabaster, AL). Tris-HCl, tris base, NaCl, NaOH, and sodium dodecyl sulfate (SDS) were bought from Sigma-Aldrich (St. Louis, MO) or Thermo Fisher Scientific, Inc. (Waltham, MA). Other chemicals and consumables were purchased from either VWR, Inc. (Radnor, PA) or Thermo Fisher Scientific, Inc. (Waltham, MA).

Ab5 (Aβ N-terminal specific antibody) and MM26-2.1.3.35.86 (Aβ C-terminal specific antibody) antibodies were a kind gift from Levites Lab at the Emory University (Atlanta, GA), and antimouse polyclonal secondary antibody was purchased from Abcam. pET-Sac Aβ(M1–42) plasmid was obtained from ADDGENE and plasmid with Aβ mutant genes was subcloned at the molecular cloning facility at Florida State University.

### 2.1. Recombinant Aβ Expression and Purification

Aβ M1–42 (M is the first methionine residue) and mutant Aβ M1–42 (R5A, H^13^H^14^AA, and K16A) plasmids was transformed and expressed in BL21(DE3) PlysS Star *Escherichia coli* cells and purified as previously reported[77,78]. Typically, 50 mL cells were grown overnight in ampicillin containing LB broth. A part of this overnight culture was inoculated to a liter of ampicillin containing LB media and the cells were allowed to grow till an optical density of 0.6-0.8 at which time they were induced with 1mM IPTG and further allowed to grow for 16 h at 37 °C. The cells were harvested by centrifugation at 10000 rcf the following day and lysed by sonication multiple times and centrifuged to acquire inclusion bodies. Inclusion bodies were then resuspended in 6 M urea, sonicated again, and centrifuged at 15000 rcf. Pellet was discarded and the supernatant was filtered with a 0.2 μm hydrophilic PVDF filter to get rid of the cell debris. Pure Aβ(M1-42) or mutant was obtained by running the filtrate through high-performance liquid chromatography (HPLC) using a Zorbax C8 column preheated at 80 °C. Purified Aβ was lyophilized using a cold lyophilizer and stored at −80 °C for future use. Finally, to obtain aggregate-free Aβ monomers, HPLC-purified Aβ (∼1 mg) was resuspended in 490 μL of sterile water and allowed to solubilize at room temperature for 30 min and 10 mM NaOH was then added to the mixture and incubated at room temperature for 10 additional minutes. The sample was then subjected to size exclusion chromatography in 20 mM tris pH 8.0 using a Superdex-75 HR 10/30 column fractionating at a flow rate of 0.5 mL/min and attached either to an AKTA FPLC system (GE Healthcare, Buckinghamshire) or a BioLogic DuoFlow system (BioRad). Monomers eluted between the fractions 24 to 28 that were collected, and the concentrations were determined using UV absorbance on a Cary 50 UV–Vis spectrometer (Agilent Technologies, Inc., Santa Clara, CA) with an extinction coefficient of ε = 1450 cm^−1^ M^−1^ at 276 nm. Matrix-assisted laser desorption ionization-time of flight (MALDI-tof) spectrometry was used to determine the purity and integrity of the peptide. Purified monomers were either placed on ice or stored at 4 °C and utilized for experiments on the same day of purification.

### 2.2. Thioflavin-T (ThT) Kinetics

Aβ monomers (25 μM) was incubated with 75, GM1, GM3 or GD3 in 20 mM Tris in the presence of 50 mM NaCl and 50 μM ThT. Fluorescence kinetics were obtained in corning 96-well plates (black) in a Biotek Synergy well plate reader at 37 °C monitored every 30 min with shaking for 10 s before every read. The fluorescence data were processed and using Origin 8.0 upon normalization as done earlier.

### 2.3. Isolation of Oligomers

Aβ oligomers generated if any, from ganglioside incubations were purified by first removing high molecular weight fibrils by centrifugation at 130[000*g* for 20 minutes at 4 °C on a Beckman Coulter ultracentrifuge using a TLA55 rotor. The soluble supernatant was then run on a Superdex-75 SEC column as described above. Each fraction obtained had a total volume of 500 µL, and oligomers were found to be in the 15 to 18^th^ fraction; their concentration was determined by UV–vis spectroscopy, as described above. Samples were either placed on ice and used for experimentation immediately or lyophilized and kept at −80 °C for extended storage prior to experimentation.

### 2.4. Electrophoresis and Immunoblotting

Samples were run in polyacrylamide gel in presence of SDS by diluting samples in 1× Laemmli loading buffer onto either 4–12% Invitrogen or 4–20% Bis-Tris BioRad TGX gels. Prestained or unstained molecular-weight markers (Novex Sharp Protein Standard, Life Technologies) were run alongside samples in the gel. Proteins separated based on molecular weight onto the gel were transferred onto a 0.2 μm western blot membrane (GE life sciences) using a thermo scientific transfer cassette for 15 min. The blot with the bound protein was heated in 1X PBS for 1 min in a microwave oven followed by blocking at 25 °C in 5% nonfat dry milk with 1% Tween 20 in PBS for 1.5 h. Subsequently, the blots were then probed at 4 °C for 16 h with Ab5 monoclonal antibody (for detecting 1-16 amino acids in Aβ) or Ab42.2 [79] (for detecting C-terminal region in Aβ42) in a 1:6000 dilution, followed by secondary antibody incubation, with a 1:6000 dilution horseradish peroxidase-conjugated anti-mouse antibody, for 1.5 h at 25 °C. The blot was then imaged using a Super Signal West Pico Chemiluminescent peroxide substrate kit (Thermo Fisher Scientific).

### 2.5. Dynamic Light Scattering analysis

DLS data were collected on a Zetasizer Nano S instrument (Malvern, Inc., Worcestershire, U.K.) upon scanning a total of 15 scans on samples (5-10 µM) equilibrated for 30 s before scans obtained for 10 s each for every sample. The data were exported using the Zetasizer software provided by the manufacturer and graphed using OriginLab 8.0.

### 2.6. MALDI-Tof mass spectrometry

Molecular weights of SEC-purified Aβ42 monomers and its mutants were confirmed by MALDI-Tof analysis by using 100-120 ng of freshly purified Aβ42 monomers (fraction 23; Figure S2 a, d, g, and j) in 20 mM Tris buffer at pH 8.0). which was spotted onto a Bruker MSP 96 MicroScout Target with a 1:1 ratio of sample:sinapinic acid matrix (saturated with acetonitrile and water) and dried. MALDI-ToF mass spectrometry was performed on a Bruker Datonics Microflex LT/SH ToF-MS system. The data obtained were processed and plotted using the Bruker flexAnalysis software (Bruker Daltonics) and Origin Pro.

## 3. Results and discussion

### 3.1.1 Carbohydrate moieties on gangliosides dictate oligomerization of Aβ

Yamamoto and colleagues demonstrated the selectivity of Aβ40 and its pathogenic variants to aggregate in the presence of various gangliosides that varied primarily in their sugar distributions[71]. We have also showed the selectivity of Aβ42 to form oligomers only in the presence of glucose-containing synthetic polymer pendants[80]. Based on these, to specifically understand how sugar distributions on ganglioside micelles effect on the temporal kinetics of Aβ42 aggregation, GM1, GD3 and GM3 were chosen based on their cell distribution and structural differences (Figures 1a, b and c). GM1 contains a branched pentasaccharide units with a 1→ 4 linked galactose and galactosamine units that is absent in GM3, while in GD3 they are replaced by two α-linked N-acetylneuraminic acids (Figures 1a, b and c).

**Figure 1:**
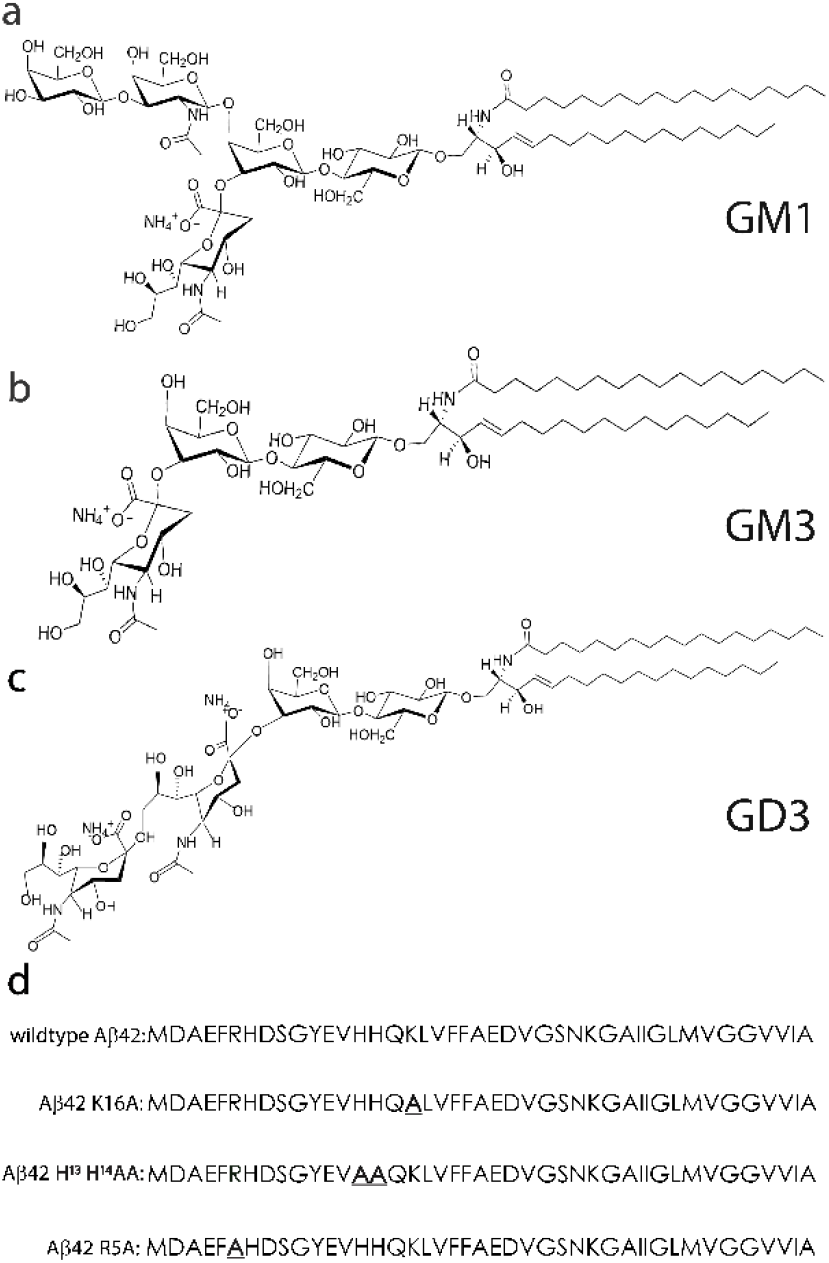
(a, b, and c) Chemical structures of GM1, GM3, and GD3 gangliosides respectively. (d) Sequences of wt-Aβ, AβK16A, AβH^13^ H^14^AA, and AβR5A.

These differences make them ideal candidates to investigate the effect of terminal sugars that are present on the surface of the micelles exposed to interaction with Aβ. To investigate this, freshly purified seed-free Aβ monomers (25 μM) buffered in 20 mM Tris (pH 8.0) containing 50 mM NaCl were incubated with 75 µM of each GM1, GD3 and GM3 respectively incubated individually at 37 °C. The concentration of 75 µM was chosen for the three gangliosides to ensure that they were above their critical micelle concentrations[81,82]. Furthermore, common sphingolipid binding domain in Aβ was determined to reside between residues 5 to 16, with R5, H13, H14 and K16 being the key players in GM1 binding[83]. Therefore, in addition to the investigation on the sugar distributions on gangliosides, to identify key amino acid residues on oligomerization of Aβ, we introduced three mutations, R5A, H^13^H^14^ AA, and K16A in our study (Figure 1 d-f).

First to see the temporal differences in aggregation due to ganglioside micelles ThT fluorescence kinetics assay was used. Wildtype Aβ (wt-Aβ) in the absence of gangliosides showed a lag time of 5-6 h (▴; Figure 2a) while the presence of all three gangliosides (GM1, GM3, or GD3) abrogated the lag times completely and thereby increased the rate of aggregation (■, •, and ..6; Figure 2a). AβR5A in the absence of lipids aggregated with an increased lag time of 8h as compared to the wildtype (▴; Figure 2b). The mutant also displayed a slower aggregation rate in the presence of GM1 and GM3 micelles with a lag time of 4 h (■, and •; Figure 2b) but GD3 micelles had no effect on the aggregation (▴; Figure 2b) suggesting that the positively charged arginine is essential to interact with GM1 and GM3 and not GD3. Removal of lysine at the 16^th^ position (AβK16A) significantly increased the lag times of both in the absence (control) and presence of lipids (Figure 2c). While the control AβK16A displayed a lag-time of >16 h (▴; Figure 2c) those incubated with gangliosides did not show well-defined lag times but rather slow rates of aggregation (■, •, and ▴; Figure 2c) suggesting that the lysine is also important for interactions with the gangliosides. To observe the effect of histidines at 13^th^ and 14^th^ positions on aggregation, AβH^13^H^14^AA double mutant was used. AβH^13^H^14^AA in the absence of lipids showed a delayed lag time of 10-12 h (▴; Figure 2d). Incubation with GM1 or GM3 gangliosides showed a lag time of ∼10 h (■, and •; Figure 2d) and with intensities of fluorescence lower than the other mutants or wildtype possibly indicating a slower and somewhat less ThT positive fibrils. However, AβH^13^H^14^AA in the presence of GD3 ganglioside micelles there was a low in fluorescence intensity but without any lag time (▴; Figure 2d). Taken together this data suggests that R5, K16, H13, and H14 residues on Aβ seem to influence the interactions with gangliosides with varying effects on aggregation.

**Figure 2:**
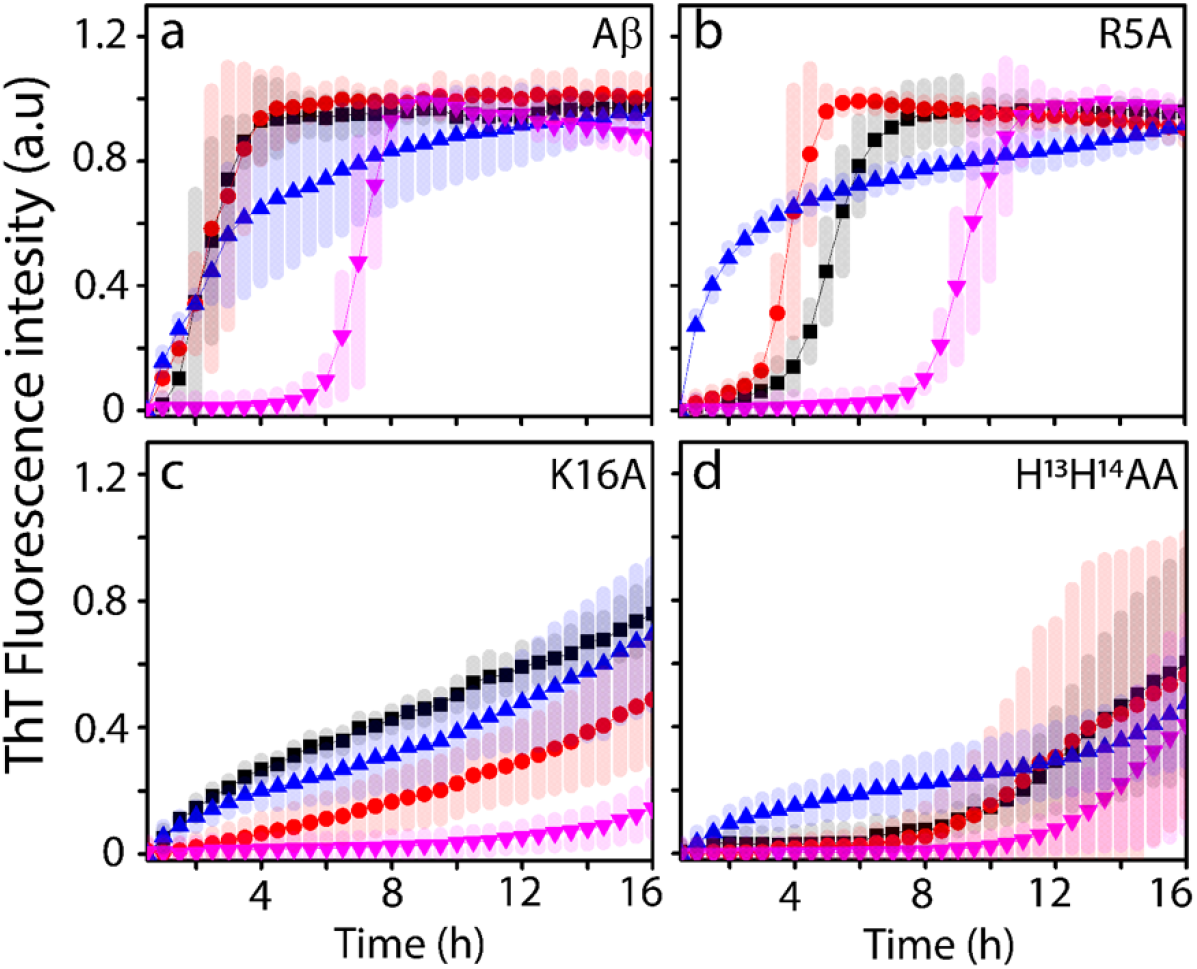
(a, b, c and d) Normalized ThT fluorescence kinetics of 25 µM wt-Aβ, R5A Aβ mutant, K16A and H^13^H^14^AA mutants of Aβ respectively without (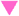; control) or with 75µM GM1 (■), GM3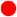;, or GD3 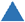; ganglioside lipid micelles buffered in 20mM tris buffer pH 8.00 with 50 mM NaCl and 50 µM ThT monitored with intermittent shaking for 16-18 h at 37°C in BioTeK synergy HTX 96-well plate reader.

Next, we sought to investigate whether the change in aggregation kinetics due to gangliosides results in the formation of stable or transient oligomeric intermediates. To do so, aliquots of the samples analyzed for Figure 2 were removed after 3, 6, and 24h, and subjected to SDS-PAGE/immunoblotting analysis. Before loading onto electrophoresis, the samples were centrifuged at 130,000 g for 30 mins to ensure that all high molecular weight species such as protofibrils and fibrils were sedimented and that only the low-molecular-weight soluble oligomers remained in the supernatant. At each time point mentioned above, supernatants of the samples were electrophoresed on a partly denaturing SDS-PAGE (without sample boiling) and immunoblotting using Aβ-specific antibodies. As expected, wt-Aβ in the absence of gangliosides (control) showed only high molecular weight fibrils that sediment upon centrifugation (lanes 4, 8, 12, 16, 20 & 24; Figure 3a). In the presence of GM1 micelles, wt-Aβ showed oligomeric bands near 40-50 kDa as well as near 15 and 4 kDa within 3h (lane 1; Figure 3a). As we have previously shown, the low molecular weight bands are likely to be formed due to dissociation in SDS during electrophoresis[26]. These oligomeric bands remained in the supernatant even after 24 h suggesting that they are thermodynamically stable (lanes 5, 13, & 21; Figure 3a). In the presence of GM3 micelles, wt-Aβ showed only trimeric and monomeric bands at 3 h (lane 2; Figure 3a) but displayed high molecular weight aggregates at 6 and 24h that sedimented upon centrifugation as the reaction progressed (lanes 10, 14, 18, & 22; Figure 2a). In case of samples incubated with GD3 micelles, wt-Aβ showed oligomeric bands ranging between 40 and 160 kDa along with trimeric and monomeric bands in addition to high molecular weight fibrils that failed to enter the gel after 3h (lane 3; Figure 3a). However, the 40 kDa band sedimented out upon centrifugation suggesting that these oligomers were likely not present in the sample but were generated due to dissociation during SDS-PAGE; however, a faint band of 160 kDa persisted suggesting that GD3 induced the formation of higher molecular weight oligomers (lanes 7, 15, &23; Figure 3a). Fibrillar bands started to emerge in all the reactions including control after 6 24 h (lane 9-12 and lane 17-20; Figure 3a). Additionally, the absence of any monomeric species in the supernatant fraction of GM3 and control reactions after 24 h suggests near-complete sedimentation of the higher molecular weight species upon centrifugation.

**Figure 3:**
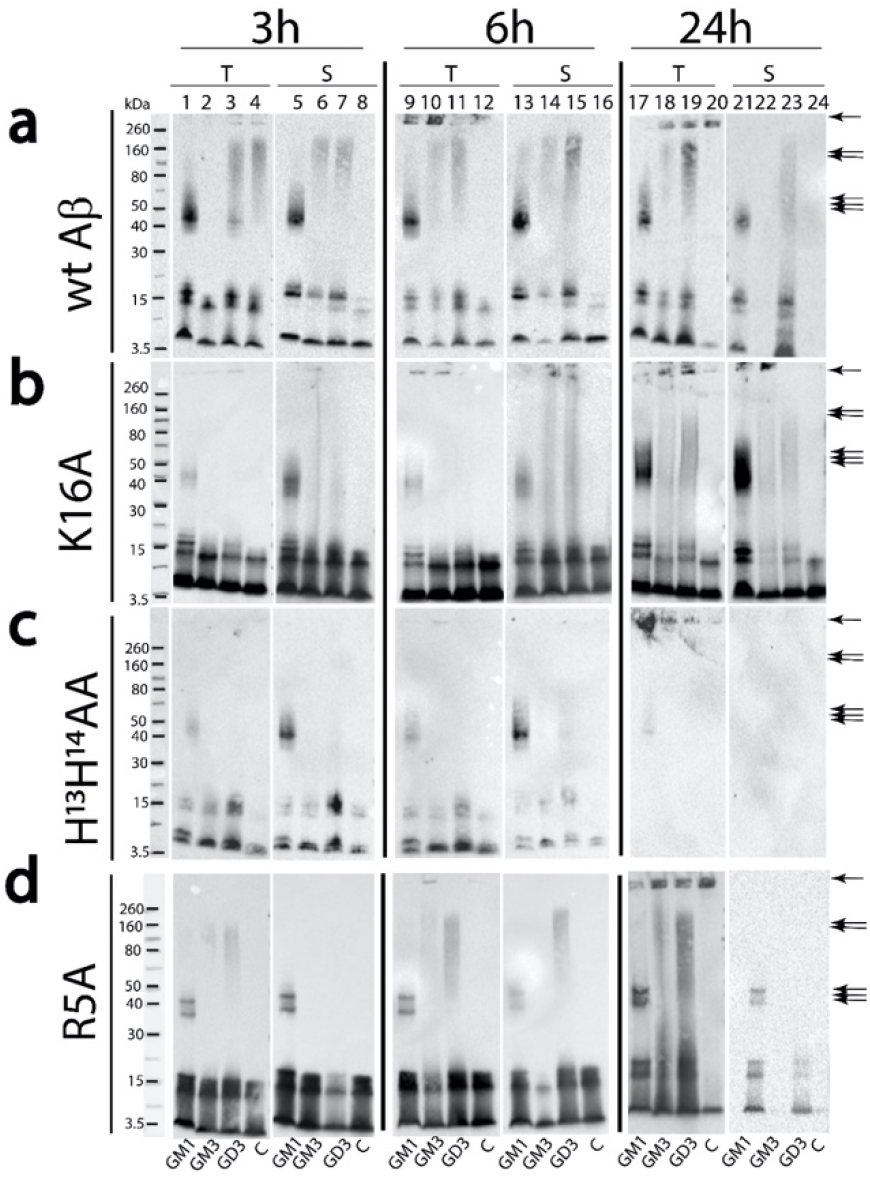
(a, b, c, and d) Partially denaturing SDS-PAGE immunoblots of 25 µM wt-Aβ, AβK16A, AβH^13^ H^14^AA, and AβR5A, respectively, in the presence of 75µM GM1, GM3, or GD3 micelles or without micelles, i.e., control (C). Samples were incubated in 20 mM Tris pH 8.00 with 50 mM NaCl at 37 °C and centrifuged at 130,000g for 30 minutes at 4°C at intervals of after 3, 6, and 24 h. The total and supernatant samples were run on gel respectively and probed with either Ab5 monoclonal antibody that binds to the N-terminal of Aβ sequence for (a), (b), and (c) or Ab42.2 monoclonal antibody with a C-terminal epitope of Aβ42 for (d). All single arrows represent fibrils, double arrows represent high molecular weight species and triple arrows represent low molecular weight oligomers.

In addition to the wildtype, immunoblots of samples with Aβ mutants in the presence of GM1, GM3, and GD3 gangliosides in similar pH and buffer conditions were analyzed. In the case of AβK16A, 40-50 kDa oligomeric bands (lane 1; Figure 3b), albeit diminished as compared to the wildtype, were observed only with GM1 after 3h and not with other lipids. However, importantly, not only did these oligomers persist even after 24h, but the intensity of the oligomeric band was also significantly enhanced with time suggesting a slower rate of formation of these oligomers (lanes 5, 13, & 21; Figure 3b), which parallels the ThT kinetic data (Figure 2c). Furthermore, trimeric and monomeric bands were only seen in all the samples consistent with the dissociation of high molecular weight species during electrophoresis. For AβH^13^H^14^AA, incubation with GM1 micelles resulted in the formation of 40-50 kDa oligomers within 3h and 6 h that persisted in the supernatant (lanes 5 & 13; Figure 3c). However, after 24 h, these oligomer bands diminished that eventually got sedimented out (lane 21; Figure 3c). No oligomers were found to form upon incubation of GM3 or GD3 (Figure 3c). In case of AβR5A, we used the C-terminal-specific monoclonal antibody Ab42.2[79] since Ab5 antibody is an N-terminal specific antibody, which could render the antibody ineffective due to R5A mutation as we discovered a low binding efficiency upon probing (Figure S1). Therefore, probing of the immunoblot with Ab42.2 antibody AβR5A mutant interestingly showed faint oligomeric bands at 50-60 kDa with GM1that persisted in the supernatant even after 24 h (lanes 5, 13, & 21; Figure 3d). Incubation of the mutant with GM3 or GD3 showed 60-260 kDa for respectively at 3 h and 6 h that sedimented out upon centrifugation (lanes 2, 3, 10, 11, 23, & 23; Figure 3d). The control in the absence of lipids showed no oligomer bands at any time. In sum, the data suggests that the two galactose units present in GM1 ganglioside seem to be critical for the oligomerization of Aβ. In Aβ, the R5 residue seems to be important for the formation of oligomers while K16 delays it. The H13 and H14 on the other hand seem to affect the stability of the oligomers formed.

### 3.1.2 Isolation of discrete GM1- and GD3-catalyzed oligomers suggests their relative stabilities

As shown in Figure 3, we observed that GM1 induces the formation of 40-50 kDa oligomers while GD3 induced the formation of dispersed higher molecular weight oligomers of wt-Aβ (40-160 kDa) while GM3 did not show oligomer formation. These oligomers were present in the soluble supernatant fraction upon ultracentrifugation and persisted for 24 h. We questioned whether the oligomers could be fractionated by SEC further providing evidence for their relative thermodynamic stabilities to withstand dissociation. Therefore, to quantitatively analyze the emergence and stability of the oligomers formed, aliquots of wt-Aβ incubated with GM1, GM3, and GD3 were taken after 3, 6, and 24h and fractionated on a Superdex-75 size exclusion column (Figure 4) after removing the high molecular weight species such as protofibrils and fibrils by centrifugation at a centrifugal force of 130,000xg. After 3h, the GM1-incubated samples showed a large oligomer peak that fractionated near the void volume followed by GD3-incubated ones (Figure 4a, b, d, and e). The GM3-incubated sample showed a negligible amount of oligomer which showed no considerable depletion over 24h (dotted line and box; Figure 4). The GM1-incubated samples showed no decrease for 6 h, and the amount of oligomers was reduced to 60% after 24 h suggesting that the oligomers are transiently stable (black line and box; Figure 4). The GD3-incubated samples showed a progressive decrease in the amount of oligomers suggesting that they are more transient than GM1 oligomers besides being less abundant.

**Figure 4:**
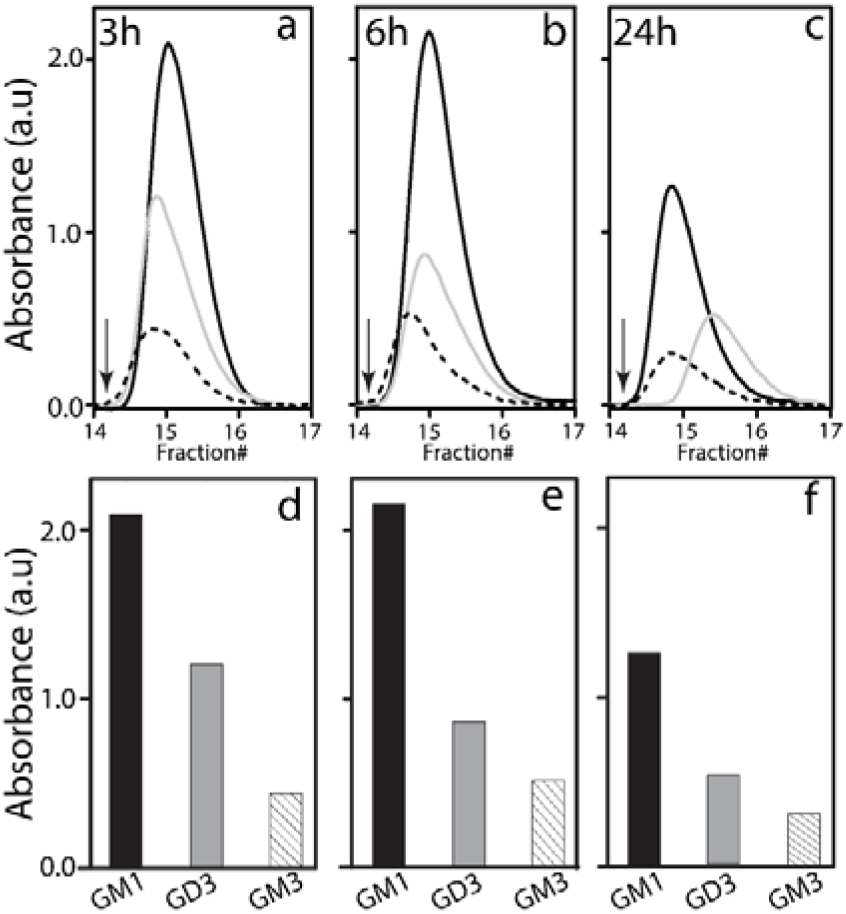
Quantitative stability analysis of oligomers by SEC fractionation. SEC chromatograms (a-c) of the supernatants of Aβ samples (25 µM) incubated in the presence of 75µM GM1 (**—**), GM3 (---), or GD3 (**—**) ganglioside micelles buffered in 20 mM Tris pH 8.0, 50 mM NaCl at 37 °C after centrifugation at 130,000xg for 30 min after 3h (a), 6h (b) and 24 h (c) of incubation. at 37°C. The arrow indicates the void volume. (d-f) Bar diagram showing the corresponding integrated area under the peak for SEC chromatograms for GM1, GD3, or GM3 incubated samples.

After analytically analyzing the oligomers, oligomers were subjected to a preparative scale for further characterization. To do so, a preparative scale SEC fractionation was performed on a Superdex-75 column. As expected, with GD3, two distinct fractionations were observed in the SEC chromatogram: one near the void volume between fractions 13 and 16 and one between 20 and 24 (Figure 5a). The former fractions typically show oligomers, and the latter fractions show monomers based on our previous observations with different lipid-micelle catalyzed oligomers[26]. It is important to note that these SEC-fractionated oligomers are free of monomers but are complexed with the lipid (∼<10%) as we showed before [77]. To deduce the molecular weight of these freshly isolated wt-Aβ oligomers, they were subjected to SDS PAGE electrophoresis followed by immunoblotting with Ab5 antibodies. A molecular weight of 60-260 kDa dispersed protein bands was observed with faint dimeric/trimeric bands as well as monomeric bands (Figure 5b) that was similar to the banding pattern observed prior to fractionation. Size analysis of GD3-derived oligomers was performed on a dynamical light scattering (DLS) which showed a monodisperse hydrodynamic diameter centered around 21 nm (Figure 5c).

**Figure 5:**
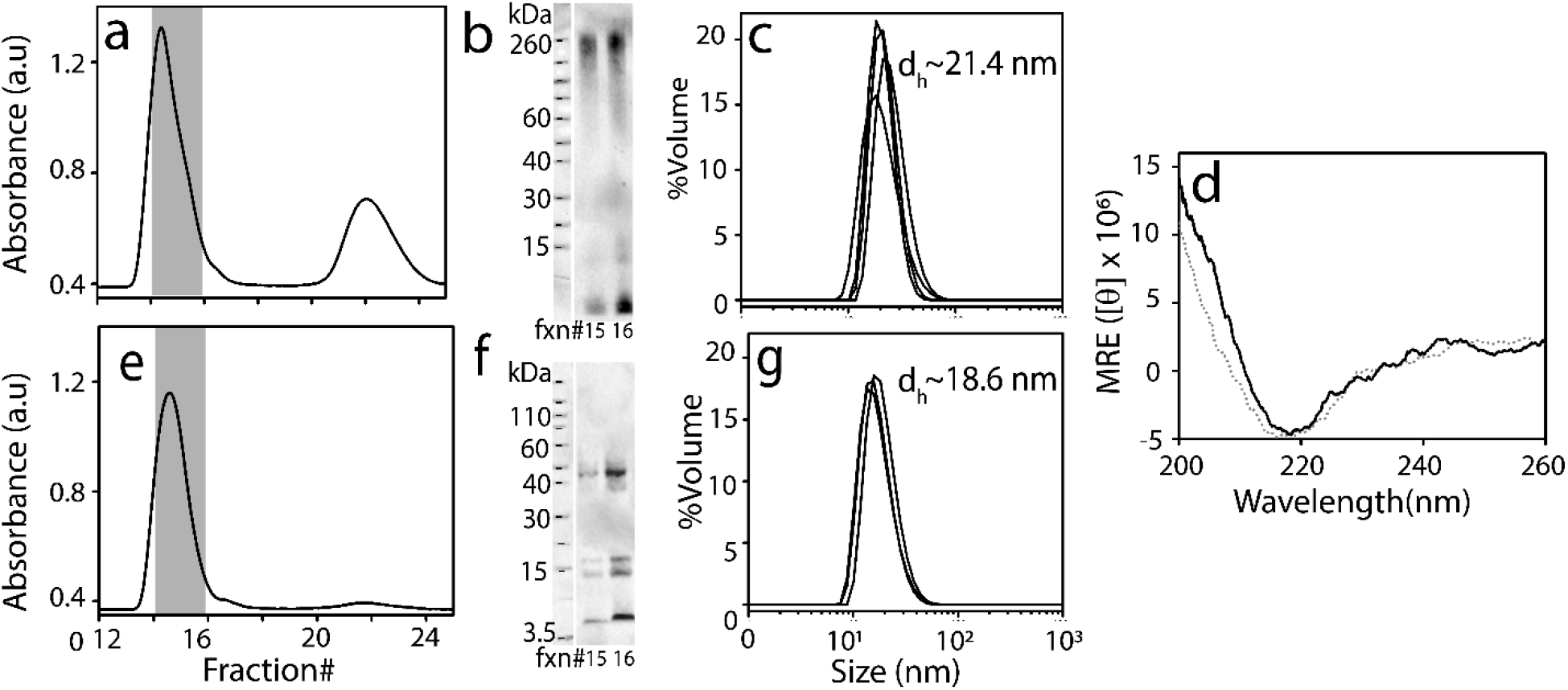
(a) SEC chromatogram for isolation of Aβ oligomers generated in the presence of 75µM GD3 ganglioside micelles buffered in tris pH-8.0 50 mM NaCl after incubation at 37 °C for 24 h and centrifugation at 130,000g for 30 min. (b) Immunoblots of SEC-isolated oligomer fraction 15–16 of Aβ oligomers generated in the presence of GD3 micelles in (a) probed by Ab5 monoclonal antibody. (c) DLS for fraction 16 of SEC-isolated Aβ oligomers (accumulation of a total of 12 scans) generated in the presence of GD3 gangliosides. (d) CD spectra (accumulation of 3 scans) of fraction 16 of SEC-fractionated Aβ oligomers generated in the presence of GD3 (**—**) or GM1(---) ganglioside micelles. (e) SEC chromatogram for isolation of Aβ oligomers generated in the presence of 75µM GM1 ganglioside micelle buffered in tris pH-8.0 with 50 mM NaCl after incubation at 37°C for 24 h and centrifugation at 130000g for 30 min. (f) Immunoblots of 15–16 from SEC-isolated Aβ oligomer fraction generated in the presence of GM1 micelles in (a) probed by Ab5 monoclonal antibody. (g) DLS for fraction 16 of SEC-isolated Aβ oligomers (accumulation of a total of 12 scans) generated in the presence of GM1 gangliosides.

The secondary structure of the isolated fractions showed a β-sheet by far-UV CD analysis with a negative minimum at 218 nm (Figure 5d). In case of the GM1-incubated sample, a similar SEC profile was observed with two fractions, one at 15-16 and the other at 22-24 although with different areas of the fractions (Figure 5e). A far decreased monomeric peak at 22-25 indicates that more oligomers or fibrils are formed with GM1 as compared to GD3. It is also likely that GD3 oligomers are unstable to undergo dissociation upon fractionation. Nevertheless, a distinct oligomeric band corresponding to 40 kDa was observed along with dimeric and monomeric Aβ bands on immunoblot of the fractions 16 and 17 (Figure 5f). Further size analysis of the oligomers with DLS showed a monodisperse hydrodynamic diameter of ∼ 15 nm (Figure 5g). Far-UV CD data showed a spectrum with a negative minimun at 218 nm indicative of a β-sheet (Figure 5d). Due to negligible yields of GM3 oligomers, if any were present, we could not obtain DLS and CD data.

## 3.2 Discussion

The interaction of Aβ with gangliosides has been well researched in the past[26,58– 60,65,67,69,77]. It has been shown that GM1, GM3, and GD3 preferentially interact with Aβ to augment its aggregation[71,72,76]. It was also demonstrated that ganglioside expression in neuronal culture is cell-type specific and that wildtype Aβ selectively binds to GM1 whereas GM3 and GD3 gangliosides have been found to modulate different variants of Aβ involved in familial AD such as Flemish, Dutch, Italian and Iowa[73]. Although the affinity of Aβ towards ganglioside interaction has been well known, what precise contributions different sugar distributions on gangliosides have towards Aβ oligomer generation and their maturation are not clearly understood. Since different gangliosides display structurally distinct sugar in their headgroups, differences in their mode of interaction with differently charged amino acid groups of Aβ is plausible. In our present work, all three gangliosides GM1, GM3, and GD3 have galactose, and N-acetylneuraminic acid joined to the common ceramide fatty acid backbone, while GM1 has an additional galactose and N-acetyl galactosamine, GM3 is devoid of these two sugars and has only three saccharide units (Figure 1). GD3, on the other hand, has two N-acetylneuraminic acid and two glucose units. These structural differences lead to the variation in overall polarity and hydrophobicity of the ganglioside surfaces, due to which its variable interaction with Aβ becomes feasible leading to the formation of different oligomeric aggregates. This is supported by our previous observation wherein the importance of stereochemistry, hydrogen bonding patterns of different sugar groups in modulating the aggregation pathway of Aβ to cause oligomerization or protofibril formation was established[80]. In our present study, we observed the formation of distinct soluble oligomer with molecular weight around 40 kDa in the presence of GM1. However, we observed the formation 110-260 kDa of high molecular weight Aβ oligomer in the presence of GD3 micelles and GM3 failed to generate any identifiable oligomer. Our parsimonious stability analysis suggested that while all oligomers are somewhat transient, GM1-induced ones are by far more abundant and stable. Interaction of Aβ with GM3 micelles led to the formation of high molecular weight aggregate with an augmented rate of aggregation. Furthermore, Williamson et al. reported that H13 as a key mediator in Aβ-GM1 binding.[84] They also suggested that the overall cationic charges on Aβ N-terminal region are crucial for the interaction between Aβ-GM1. This is consistent with our finding that H13 and H14 are the most important residues for oligomer formation and maturity.

Mutation of K16 seems to delay the formation of oligomers with GM1 but stabilizes them while mutation of R5 does not affect GM1 oligomer formation but only diminished band intensities of the oligomers formed. These data indicate that specifically the cationic residues in the region between 12-16 in Aβ specifically are involved in the interactions between the galactose and/or N-acetyl galactosamine.

However, almost all the cationic residues on Aβ investigated, i.e., R5, H13H14, and K16 seem to have a significant effect on oligomer formation with GM3 and GD3. Nonetheless, our findings suggest that both K16 and H13H14 are crucial for GM1 interactions and are important residues for oligomer formation and maturation. This conclusion also parallels our previous investigation on anionic fatty acid micelle-induced induced 12mers of Aβ, whose ring-like structure was held together by a key salt-bridge involving K16 and protonation states of the histidines [85]. Overall, our work indicates that the oligomerization of Aβ in presence of gangliosides is mainly guided by the interactions with the sugar distributions on the headgroups, and positively charged amino acid residues on the polar N-terminal end of Aβ contribute greatly to these interactions.

## 4. Conclusions

The results presented in this study delineate the significance of different sugar moieties attached to lipid chains, specifically gangliosides, and the importance of amino acid sequence of Aβ in the generation of oligomers with distinct biophysical and biochemical properties. Findings from this report indicate that formation of biophysically and biochemically distinct Aβ oligomers in presence of ganglioside is highly dependent on the type of sugar groups present on them and this mechanism involves interaction with key amino acid residues on the Aβ sequence. The data also indicate that the size and structure of Aβ aggregates vary in the presence of ganglioside with different sugars. Furthermore, it indicates the importance of arginine and lysine residues as well as histidine residues near the N-terminal of Aβ conferring the formation and stability of the oligomerization. These findings may portray the molecular basis of Aβ oligomerization and propagation to trigger cytotoxicity well before the formation high-molecular-weight fibrils, and thus, significant in understanding the molecular basis for low molecular weight oligomer formation.

## CRediT authorship contribution statement

VR conceptualized and oversaw the research along with intellectual contributions from JS. JS, BJF, and SB performed all the experiments and collected data. JS and VR wrote and edited the manuscript.

## Supplementary figures

**Figure S1:**
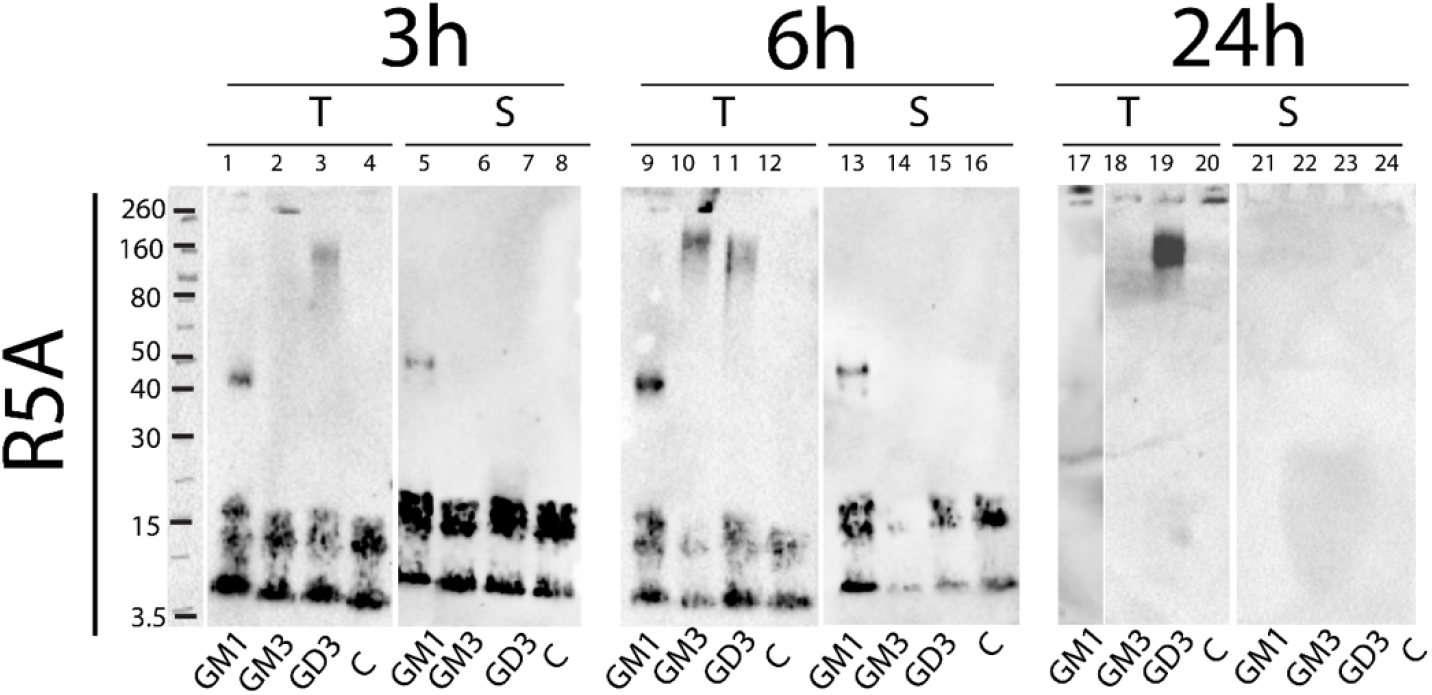
Partially denaturing SDS-PAGE immunoblots of 25 µM R5A Aβ mutant in presence of 75µM GM1, GM3, or GD3 micelles or without micelles, i.e., control (C). Samples were incubated in 20 mM Tris pH 8.00 with 50 mM NaCl and after 3, 6, and 24 h centrifuged at 130, 000g for 30 minutes at 4°C at intervals of and the total and supernatant samples were run on gel respectively and probed with Ab5 monoclonal antibody that will bind to the N-terminal of Aβ42 sequence.

**Figure S2:**
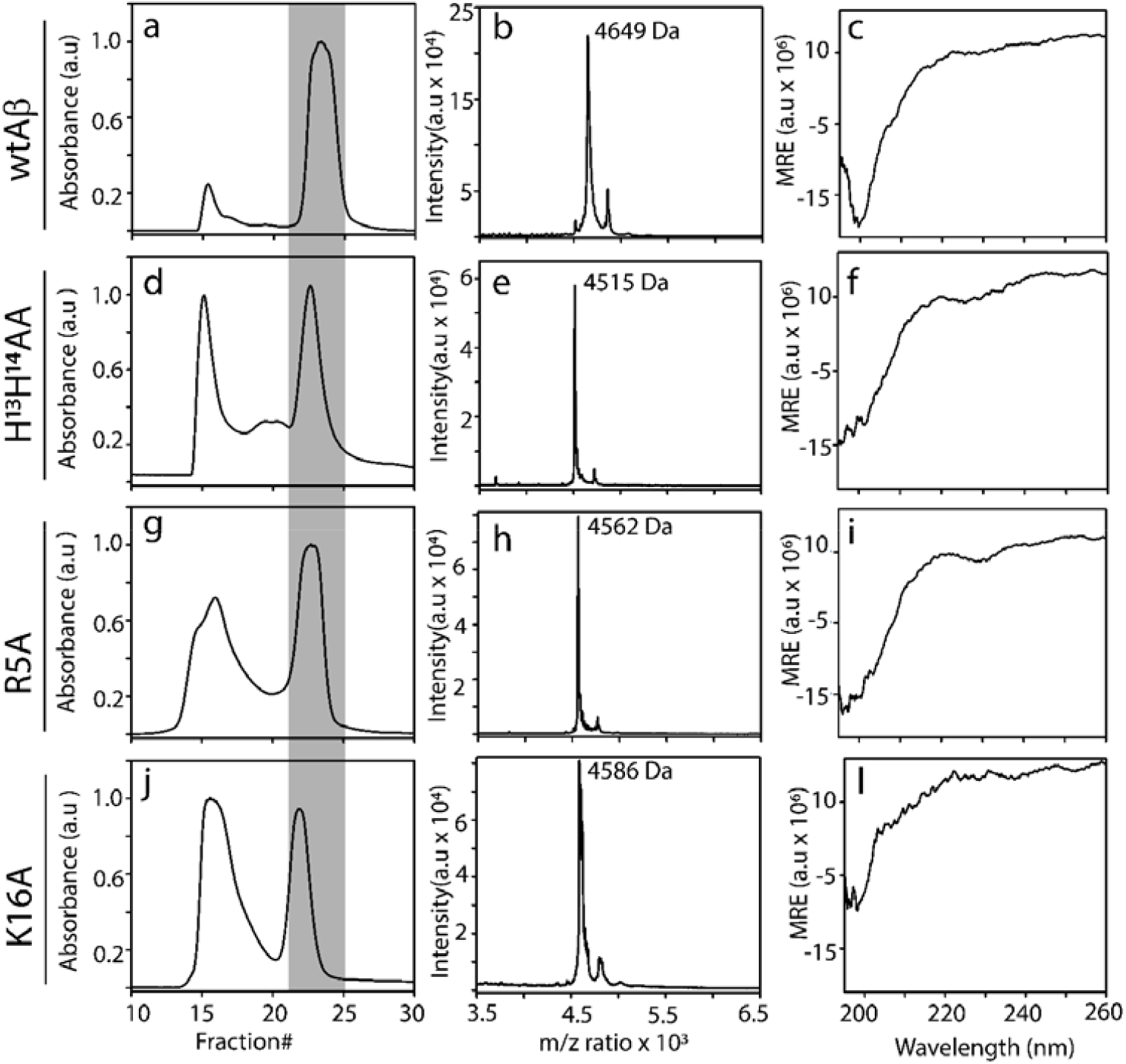
(a,d,g, and j) Size exclusion chromatogram (SEC) of wtAβ, H^13^H^14^AA, R5A, and K16A monomers respectively, after the addition of 20 mM NaOH and incubation for 15 minutes at room temperature. (b-h) MALDI-ToF mass spectrum of fraction #23 of freshly purified monomers with SEC of wtAβ, H^13^H^14^AA, R5A, and K16A respectively. (c, f, i, and l) Far-UV CD spectra fraction#23 from freshly purified 6-10 µM monomers from SEC of wtAβ, H^13^H^14^AA, R5A, and K16A respectively

